# Profiling grapevine trunk pathogens *in planta*: A case for community-targeted DNA metabarcoding

**DOI:** 10.1101/409169

**Authors:** Abraham Morales-Cruz, Rosa Figueroa-Balderas, Jadran F. García, Eric Tran, Philippe E. Rolshausen, Kendra Baumgartner, Dario Cantu

**Author notes:** Corresponding author: Dario Cantu, One Shields Ave, Davis CA 95616, USA.

## Abstract

DNA metabarcoding, commonly used in exploratory microbial ecology studies, is a promising method for the simultaneous *in planta-detection* of multiple pathogens associated with disease complexes, such as the grapevine trunk diseases. Their detection is particularly challenging, due to the presence within an individual wood lesion of multiple co-infecting trunk pathogens and other wood-colonizing fungi, which span a broad range of taxa in the Fungal Kingdom. As such, we designed metabarcoding primers, using as template the ribosomal internal transcribed spacer of grapevine trunk-associated Ascomycete fungi (GTAA) and compared them to two universal primer widely used in microbial ecology. We first performed in *silico* simulations and then tested the primers by high-throughput amplicon sequencing of (i) multiple combinations of mock communities, (ii) time-course experiments with controlled inoculations, and (iii) diseased field samples from vineyards under natural levels of infection. All analyses showed that GTAA had greater affinity and sensitivity, compared to those of the universal primers. Importantly, with GTAA, profiling of mock communities and comparisons with shotgun-sequencing metagenomics of field samples gave an accurate representation of genera of important trunk pathogens, namely *Phaeomoniella, Phaeoacremonium*, and *Eutypa*, the abundances of which were greatly over- or under-estimated with universal primers. Overall, our findings not only demonstrate that DNA metabarcoding gives qualitatively and quantitatively accurate results when applied to grapevine trunk diseases, but also that primer customization and testing are crucial to ensure the validity of DNA metabarcoding results.

## INTRODUCTION

Grapevine trunk diseases affect the longevity and productivity of grapevines *(Vitis vinifera)* in all major growing regions of the world [1-4]. They are caused by numerous species of fungi that infect and damage the wood, causing chronic infections [5-7]. Among the most common grapevine trunk diseases are Eutypa dieback (primarily caused by *Eutypa lata)*, Esca (primarily caused by *Phaeoacremonium minimum, Phaeomoniella chlamydospora*, and *Fomitiporia* spp.), Botryosphaeria dieback (primarily caused by *Neofusicoccum parvum, Diplodia seriata*, among other fungi in the Botryosphaeriaceae family), Phomopsis dieback (primarily caused by *Diaporthe ampelina)*, and Black foot (caused by *Cylindrocarpon, Campylocarpon*, and *Ilyonectria* spp.) [4, 8-11]. Because of the characteristic mixed infections, trunk diseases represent a disease complex <u>[12, 13]</u>. In addition to infections of pruning wounds by airborne and splash-dispersed spores, trunk pathogens may be introduced to a healthy vineyard by asymptomatic propagation material. Fungi associated with grapevine trunk diseases have been found in rootstock mother-plants, rooted rootstock cuttings, bench-grafts, and young grafted vines <u>[14-16]</u>. The presence of multiple species in the same vine complicates disease diagnosis and, consequently, proper timing of practices to limit infection in the vineyard and to propagate clean nursery stock.

Taxonomic identification of fungi associated with grapevine wood is currently done by the following steps: (i) plating grapevine woody tissue on nutrient-rich agar plates, (ii) hyphal-tip colony isolation to pure cultures, (iii) DNA extraction from fungal mycelium, (iv) PCR amplification of taxonomically informative loci, such as the nuclear ribosomal internal transcribed spacer (ITS), elongation factor, and β-tubulin, and (v) comparisons of amplicon sequences with sequence databases <u>[17-19]</u>. PCR-based diagnostics represent a significant improvement compared to traditional approaches that depend on morphological features for species identification and, thus, require skilled expertise in mycology [20]. However, these approaches still require an initial culturing step, which may limit the detection of slow-growing fungi. Alternatively, with species or genus-specific markers, PCR could be used to determine *in planta* the presence of certain species, thereby skipping the culturing step [21, 22]. One limitation of this approach, however, is that it may not detect all trunk pathogens in a given sample [23, 24]. Indeed, certain combinations of fungi may be important in the severity of symptom expression [25]. Because trunk pathogens cause mixed infections, attempts have been made to characterize the composition of the trunk-pathogen community. For example, finger-printing techniques like Automated Ribosomal Intergenic Spacer Analysis (ARISA) [26] and Single-Strand Conformation Polymorphism (SSCP) [27, 28] have been used to compare fungal communities among different samples of grapevine wood, although these do not identify trunk pathogens to the species level. A DNA macroarray system, based on reverse dot-blot hybridization containing oligonucleotides complementary to portions of the β-tubulin locus, was developed for species-level identification, specifically for detection of trunk pathogens that cause Young vine decline [23]. We previously described a strategy, based on untargeted shotgun sequencing of metagenomic DNA and RNA, to detect and quantify trunk pathogens *in planta* simultaneously [13]. Despite clear advantages over other approaches, this method still has its limitations, such as relying on assembled genomes, as well as costly library preparation and computationally intensive analyses.

DNA metabarcoding, which has been used extensively for the analysis of microbial communities <u>[29-32]</u>, may provide a cheaper and more scalable method for the characterization of trunk-pathogen communities. This approach has already been applied to other pathosystems to address a variety of research objectives. For example, DNA metabarcoding has been used to identify candidate pathogens [33, 34] and potential biocontrol agents [35], to profile putative plant pathogens associated with insects [36], and to diagnose quarantine pathogens as part of national plant-protection programs <u>[37-39]</u>. DNA metabarcoding infers taxonomic composition of complex biological samples by amplifying, sequencing, and analyzing target genomic regions [40, 41]. The ribosomal ITS, which is under low evolutionary pressure and, thus, presents high levels of variation between closely related species, has been commonly used as a barcode for the analysis of fungal biodiversity [42, 43]. ITS is typically amplified by universal primers that anneal to the conserved flanking sequences. The “universality” of the primers, which derives from their ability to amplify a broad range of taxonomically unrelated species across the Fungal Kingdom [44], is exploited in studies that aim to profile fungal communities, typically in exploratory analyses of environmental samples. We hypothesized that although universal primers may capture broad biodiversity in exploratory analyses, they may provide less accurate representation of microbial pathogen communities than primers that are designed and optimized to amplify species known to be associated with those communities, based on prior knowledge of disease etiology. After all, grapevine-trunk diseases are one of the most widely studied disease complexes, in terms of species composition (Lamichane and Venturi, 2015). In this work, we designed and evaluated metabarcoding primers that were optimized to amplify the ITS regions of grapevine trunk pathogens. By a combination of *in silico* simulations, and analyses of ‘mock’ communities, samples from controlled inoculations, and samples from symptomatic vineyards, we demonstrated that community-customized metabarcoding provides greater more qualitatively and quantitevely accurate representation of trunk-pathogen communities than common universal primers.

## RESULTS

### Primer design, selection, and validation with target species

We designed multiple degenerate primers that target the internal transcribed spacer (ITS) of grapevine trunk-associated ascomycetes (GTAA) using the TrunkDiseaseID as reference database <u>[20]</u>. Primer potential was determined *in silico*, considering the amplicon size and estimating the number of sequence hits to the database, their alignment mismatches, and gap scores. From a total of twenty forward and three reverse degenerate primers, primers GTAA182f and GTAA526r (GTAA, hereinafter) performed the best and were selected for further testing. The GTAA primers target the entire ITS2 region with the forward and reverse primers aligning to the 5.8S ribosomal RNA and the large subunit ribosomal ribonucleic acid (LSU), respectively (Table 1 and Figure 1A). The primers produced amplicons of approximately 350 bp from isolates of seven trunk pathogens, as expected based on the amplicon size predicted from the 213 ITS sequences of ascomycetes in the TrunkDiseaseID database (301.72 ± 7.53 bp; Figure 1B). We obtained a similar amplicon size when the GTAA primers were used to amplify total DNA extracted from naturally infected grapevines with trunk-disease symptoms (Figure 1C). Failure to amplify DNA of two negative controls, grape leaves, and of the bacterium *Agrobacterium tumefaciens*, supports their specificity to Fungi (Figure 1B). Amplicon sequences matched the correct species, when aligned to the NCBI non-redundant nucleotide database using BLASTn, thereby confirming the ITS region amplified by the GTAA primers is informative for taxonomic assignments (Additional File 1).

**Table 1.**
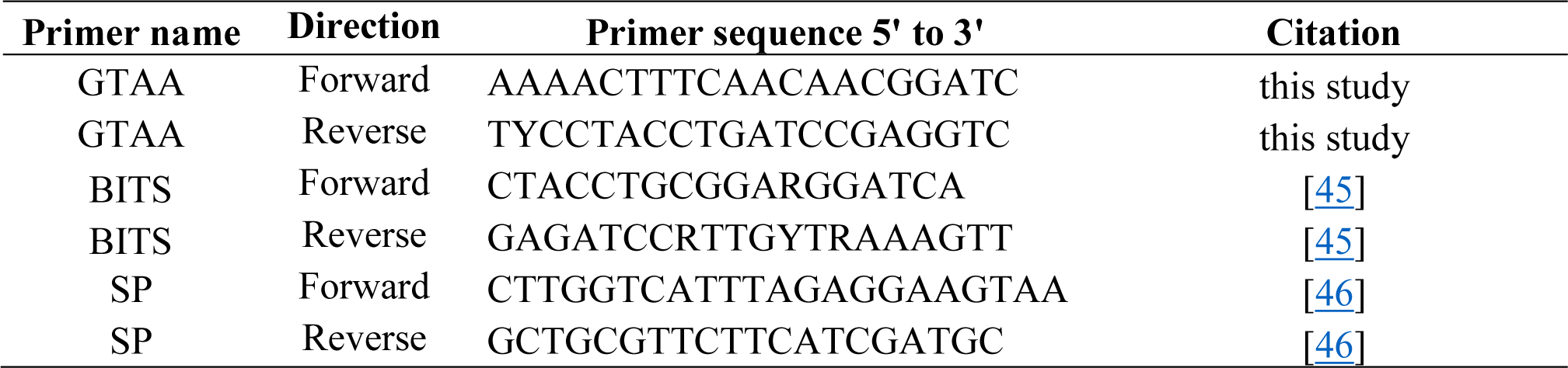
Metabarcoding primer sequences targeting the ITS region used in this study

**Figure 1.**
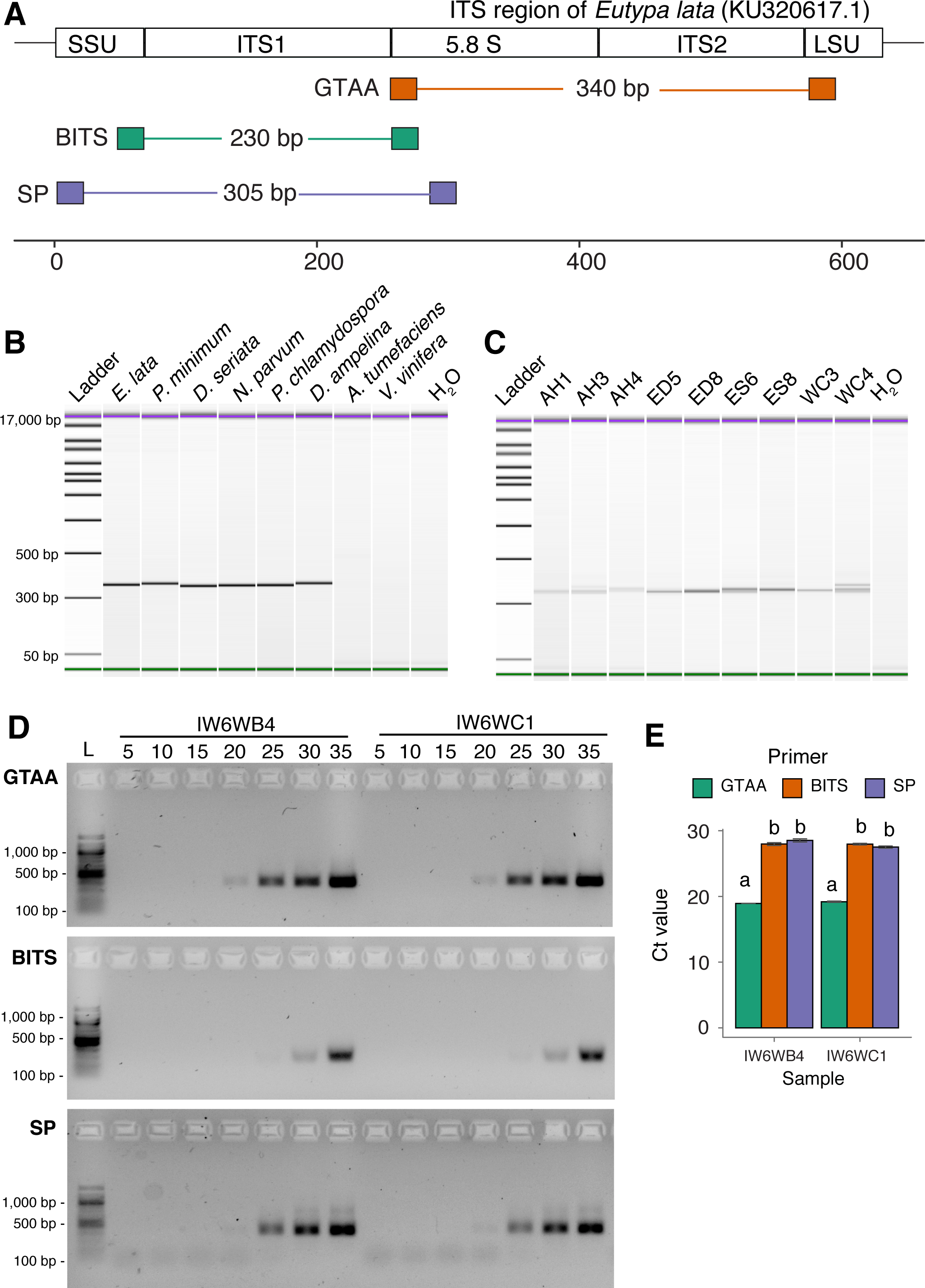
Primer design and testing. (A) Schematic representation of the annealing sites of forward and reverse GTAA, BITS, and SP primers in the fungal ribosal ITS. Reported amplicon sizes were calculated based on the ITS sequence of *Eutypa lata* (KU320617.1). (A) & (B): Bioanalyzer electropherograms showing amplicons generated using GTAA primers and (A) DNA from purified organisms as template and (C) field samples with different trunk disease symptoms. *Agrobacterium tumefaciens, V. vinifera* and nuclease-free water were included as controls. AH: apparently healthy, ED: Eutypa Dieback, ES: Esca and WC: wood cankers (no leaf symptoms). (D) PCR products from grapevines inoculated with *N. parvum* at six weeks post-inoculation, which were visible on agarose gels at five-cycle intervals, when amplified with primers GTAA, BITS, or SP. L: 100 bp Ladder. (E) Cycle thresholds (Ct) measured by qPCR of the same reactions shown in (D).

GTAA primer precision, sensitivity, efficiency, and usefulness for metabarcoding of grapevine trunk pathogens were compared to those of the BITS <u>[45]</u> and SP primers <u>[46]</u>. The BITS primers are widely used for fungal metabarcoding analysis in vineyards and grape must (e.g.: [<u>47-51</u>]), whereas the SP primers <u>[46]</u> were recently used by <u>[52]</u> for fungal microbial ecology, and implemented in the Earth Microbiome Project (http://www.earthmicrobiome.org; **Table 1 and Figure 1A**). Samples were from DNA extracted from potted grapevines either inoculated with *N. parvum* or from non-inoculated controls. By sampling PCR reactions every five cycles, the GTAA amplicon was visible on an agarose gel starting at 20 cycles, whereas those of SP and BITS were visible at 25 and 30 cycles, respectively (**Figure 1D**). Furthermore, SP produced multiple bands, which may be due to non-specific binding and/or chimeric amplicons. Based on qPCR with the same samples, the average Ct values for GTAA were approximately nine cycles lower than those of BITS (*P* < 1.85e^-04^) and SP (*P* < 3.50e^-04^) (**Figure 1E**). Overall, our findings suggest a higher affinity of the GTAA primers, when amplifying samples containing grapevine trunk pathogens.

### In *silico* simulation of amplification and taxonomic identification

We then carried out an *in silico* simulation that compared the potential amplification bias and taxonomic usefulness of GTAA, BITS, and SP primers using a comprehensive dataset of fungi associated with trunk diseases. We compiled a custom database of 521 full-length ITS sequences across 17 genera (**Figure 2A, Additional File 2**). We included only full-length ITS sequences to be able to compare primers that amplify different regions of the ITS (**Figure 1A**). *In silico* amplification of each sequence in the custom database was carried out considering all alternative sequences of degenerated primers and allowing a series of mismatches between primer and template sequences. *In silico* amplification was carried out testing all possible combinations of allowed mismatches, from 0 to 5 mismatches in the first five nucleotides of the primer (head) and 0 to 2 mismatches in the last two nucleotides of the primer (tail). GTAA primers amplified a higher number of sequences than BITS and SP primers, for every parameter tested (Figure 2B). When no mismatches between primer and target were allowed, GTAA primers amplified 85.80%, SP primers amplified 13.63%, and BITS primers were predicted to amplify none of the sequences in the database. When at least two mismatches were allowed in the tail of the primer, BITS and SP primers amplified only 16.70% and 30.33% of target sequences, respectively, whereas GTAA primers amplified 86.75%. With the most permissive parameters, GTAA primers amplified 98.08% of the sequences, and BITS and SP primers amplified 97.89% and 25.91%, respectively. The requirement of multiple mismatches for BITS primers to achieve a similar number of sequences as GTAA primers is consistent with the cycle-sampling results (Figure 1D), and suggests that GTAA primers are more efficient than BITS at amplifying the ITS of grapevine trunk pathogens.

**Figure 2.**
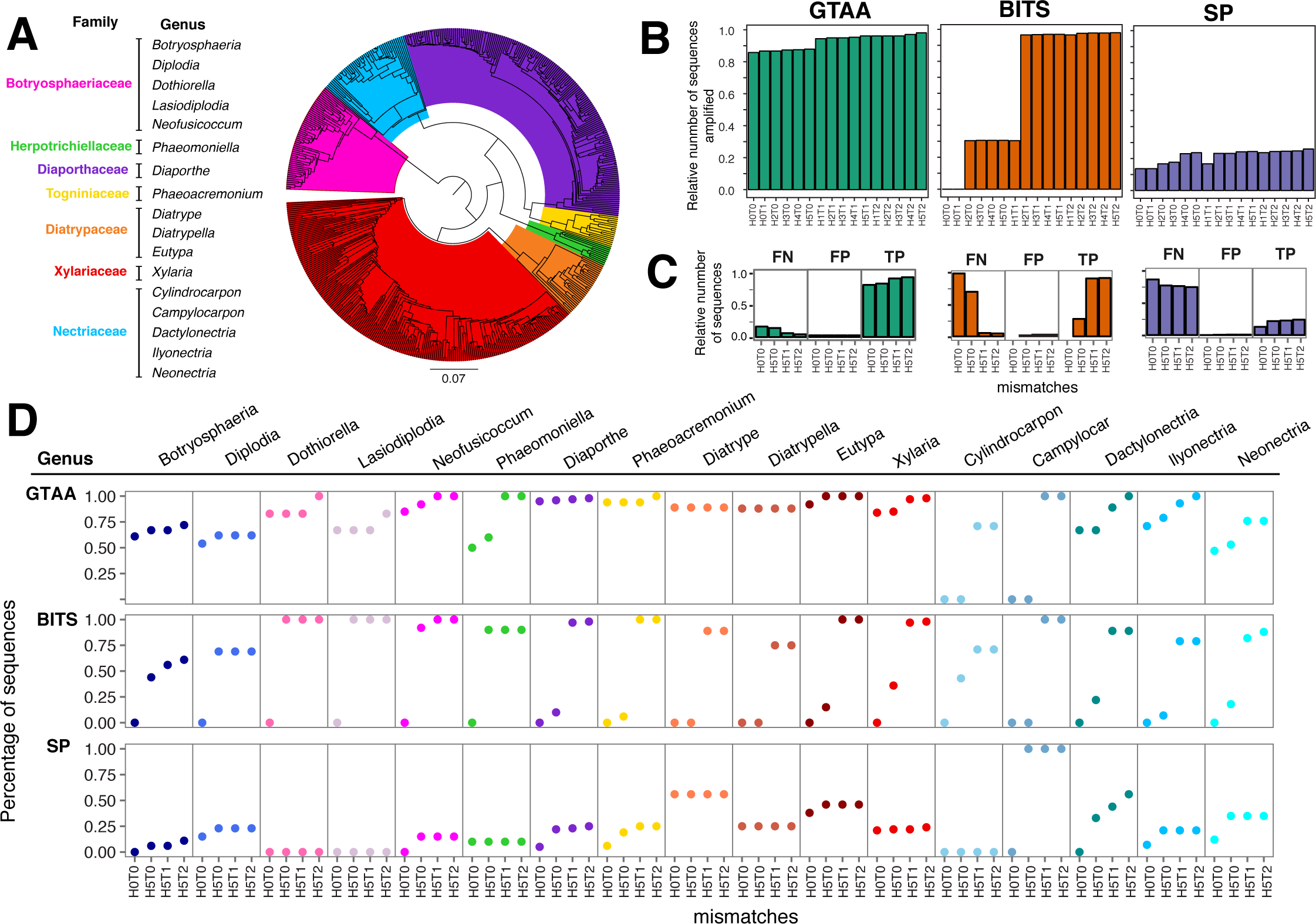
In *silico* simulation of amplification and taxonomic assignment. (A) Neighbor-joining tree of the full-length ITS sequences included in the custom database used in the simulation. (B) Barplots showing the number of sequences predicted to be amplified by each primer set at different combinations of mismatches. H: primer head. T: primer tail. Numbers correspond to the number of mismatches either in H or T. (C) Barplots showing the number of false negative (FN), false positive (FP), and true positive (TP) sequences with each primer set. (D) Percentage of sequences per genus correctly assigned with each primer set at the different mismatch combinations.

To determine if amplicons generated by GTAA primers are informative for taxonomic assignment, we analyzed with Mothur [53] the amplicons that were generated by the simulation. By comparing the assigned genera (observed) with the expected genera for each primer set we assessed false positive (FP; i.e, erroneously assigned), false negative (FN; i.e, not amplified or not assigned), and true positive (TP; i.e, correctly assigned, **Figure 2C**) rates. GTAA primers had the highest sensitivity (TP/(TP/FN)*100 = 89.50 ± 6.45%), followed by BITS (54.25 ± 47.86%), and SP (20.50 ± 2.53%). SP and GTAA primers displayed similar precision (SP: TP/(TP+FP)*100 = 97.50 ± 1.00%; GTAA: 97.00 ± 0.00%), which was higher than that of BITS primers (72.25 ± 48.18%). The different performance of the three primer sets in the simulation appeared to be mostly due to amplification bias against certain genera (**Figure 2D**). GTAA primers amplified and correctly assigned to the proper genera a larger fraction of sequences than the other two primer sets for 14 out of 17 genera tested. This was the case for the following widely distributed trunk pathogens: *Eutypa* (GTAA: 98.0 ± 4.0%, BITS: 53.8 ± 53.8%, and SP: 44.0±4.0%), *Diaporthe* (GTAA: 96.5 ± 1.3%, BITS: 51.3 ± 53.6%, and SP: 18.7 ± 9.3%), and *Phaeoacremonium* (GTAA: 95.5 + 3.0%, BITS: 51.5 + 56.1%, and SP: 18.7 ± 9.0%). BITS primers correctly assigned more sequences for *Lasiodiplodia* (GTAA: 71.0 ± 8.0%, BITS: 75.0 ± 50.0%, and SP: 0.0 ± 0.0%) and *Cylindrocarpon* (GTAA: 35.5 ± 41.0%, BITS: 46.3 ± 33.54%, and SP: 0.0±0.0%). SP primers correctly assigned more sequences for *Campylocarpon* (GTAA: 50.0 ± 57.7%, BITS: 50.0 ± 57.7%, and SP: 75.0 ± 50.0%). Overall, this simulation predicted that, unlike the two universal primer sets, GTAA primers amplify ITS of more trunk pathogens and allow taxonomic assignment with greater sensitivity (i.e., higher true positive rate) and specificity (i.e., lower false negative rate). SP primers were not included in further experiments, due to their poor performance in these early stages.

### Analysis of mock communities and infection time course

To evaluate the primers for characterizing the species composition of mixed infections, we first started by sequencing with an Illumina MiSeq and analyzing mock communities (**Figure 3A**). Although DNA was extracted from stems with no symptoms of trunk disease to be used as a pure source of grape DNA, both primer sets detected fungi, mostly belonging to the genera *Campylocarpon* and *Phaeoacremonium* (**Figure 3A**). When grape DNA was mixed with DNAs of *Pheaoa. minimum* and *Phaeom. chlamydospora* both primer sets identified the correct taxa, with small relative deviation from expected values (GTAA δ = 11.02 ± 7.0 %; BITS δ = 16.68 ± 11.39%). For mock communities including *Eutypa*, GTAA primers detected this trunk pathogen in similar amounts to the expected abundance (δ = 9.74 ± 1.10%), whereas BITS primers greatly underestimated its abundance (δ = 88.87 ± 1.27%). In mock communities with equal concentrations of DNA from *E. lata, Phaeoa. minimum, Phaeom. chlamydospora, N. parvum, D. seriata*, and *D. ampelina*, there was underrepresentation of *Eutypa* (δ = 16.70 ± 0.12%), and *Phaeoacremonium* (δ = 13.10 ± 0.63%), and overrepresentation of *Phaeomoniella* (δ = 24.34 ± 1.56%) by BITS primers (**Figure 3A**). The correlation of expected and observed abundances in these mock communities was greater for GTAA (R = 0.92) than BITS (R = 0.67; **Figure 3B**). Because DNA was mixed in equal amounts, the expected relative abundance of each genus was 16.6%. GTAA primers detected *Eutypa* at 16.42 ± 2.41%, whereas BITS primers detected this trunk pathogen at 0.05 ± 0.03%. In the case of *Diplodia*, GTAA primers estimated the abundance of the genus at 3.13 ± 0.48% and BITS primers at 28.9 ± 0.8%. Interestingly, neither primer set was able to detect properly *Diaporthe*, reporting only 0.58 ± 0.23% and 0.87 ± 0.15% for GTAA and BITS primers, respectively. Nonetheless, GTAA primers provide a better qualitative and quantitative representation of important trunk pathogens.

**Figure 3.**
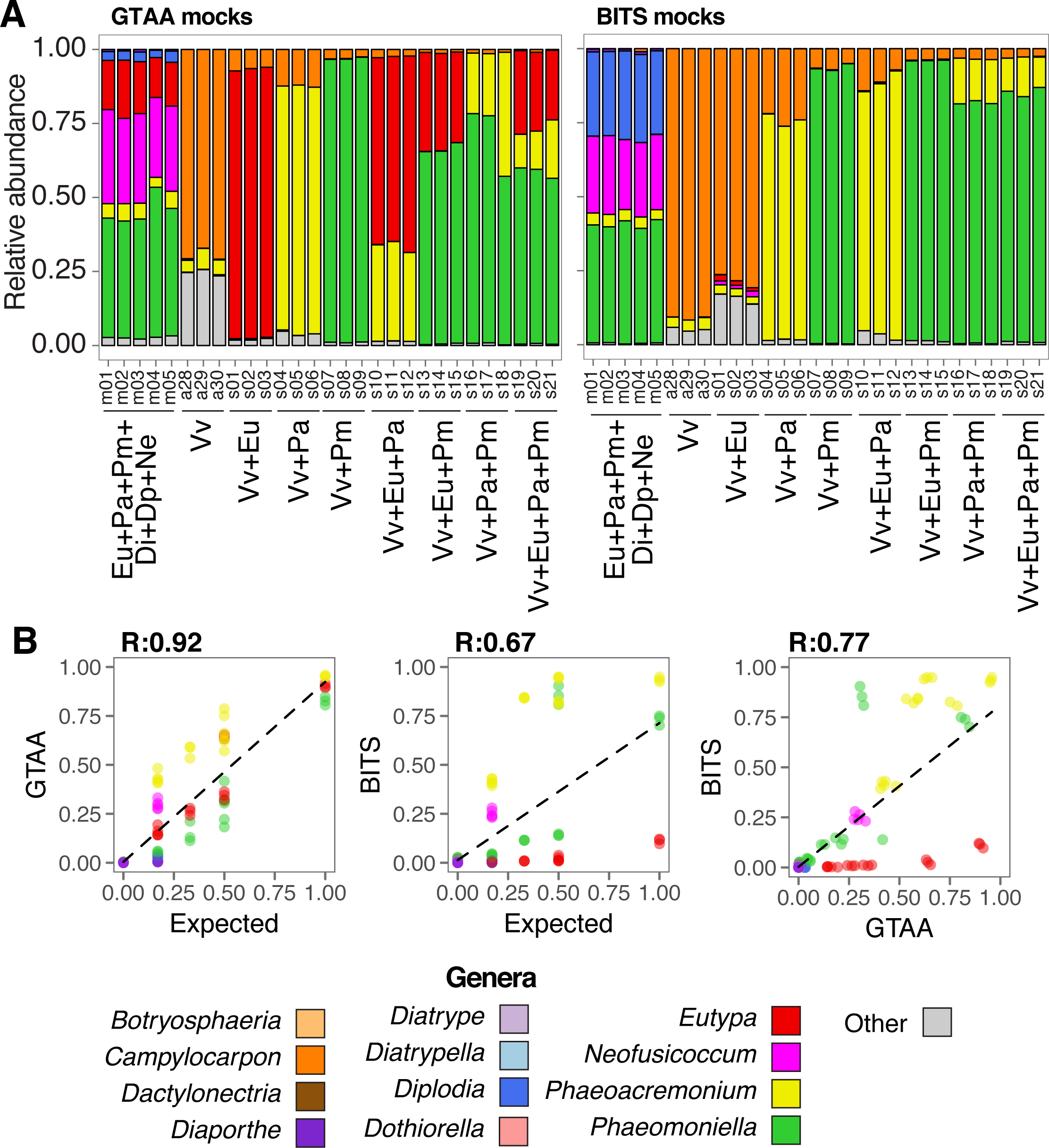
Results of DNA metabarcoding of mock communities. (A) Stacked barplots showing the relative abundance of genera in the mock communities identified using the GTAA and BITS primers. Eu: *Eutypa*, Pa: *Phaeoacremonium*, Pm: *Phaeomoniella*, Di: *Diaporthe*, Dp: *Diplodia*, Np: *Neofusicoccum*, and Vv: *Vitis vinifera.* (B) Linear correlations between observed and expected abundances for each genus contained in the mock community.

We then tested the two primers using grape samples collected at different time points after controlled inoculation with a trunk pathogen. The objective of this analysis was to determine if the metabarcoding approach could detect quantitative differences between samples at early and late stages of infection. Vines were inoculated with *N. parvum* and stem samples were collected at 24 hours, 2 weeks, and 6 weeks post-inoculation. Plants non-inoculated wounded (NIW) and noninoculated non-wounded (NINW) were included as controls. As expected, *Neofusicoccum* was predominant in the inoculated wounded (IW) samples, but absent from the controls (**Figure 4**), except for a single NIW sample, possibly due to cross-contamination during wounding or from contamination of the propagation material. Both primer sets revealed a five-fold increase in the average percentage of *Neofusicocum* between 24 hours and 6 weeks post-inoculation.

**Figure 4.**
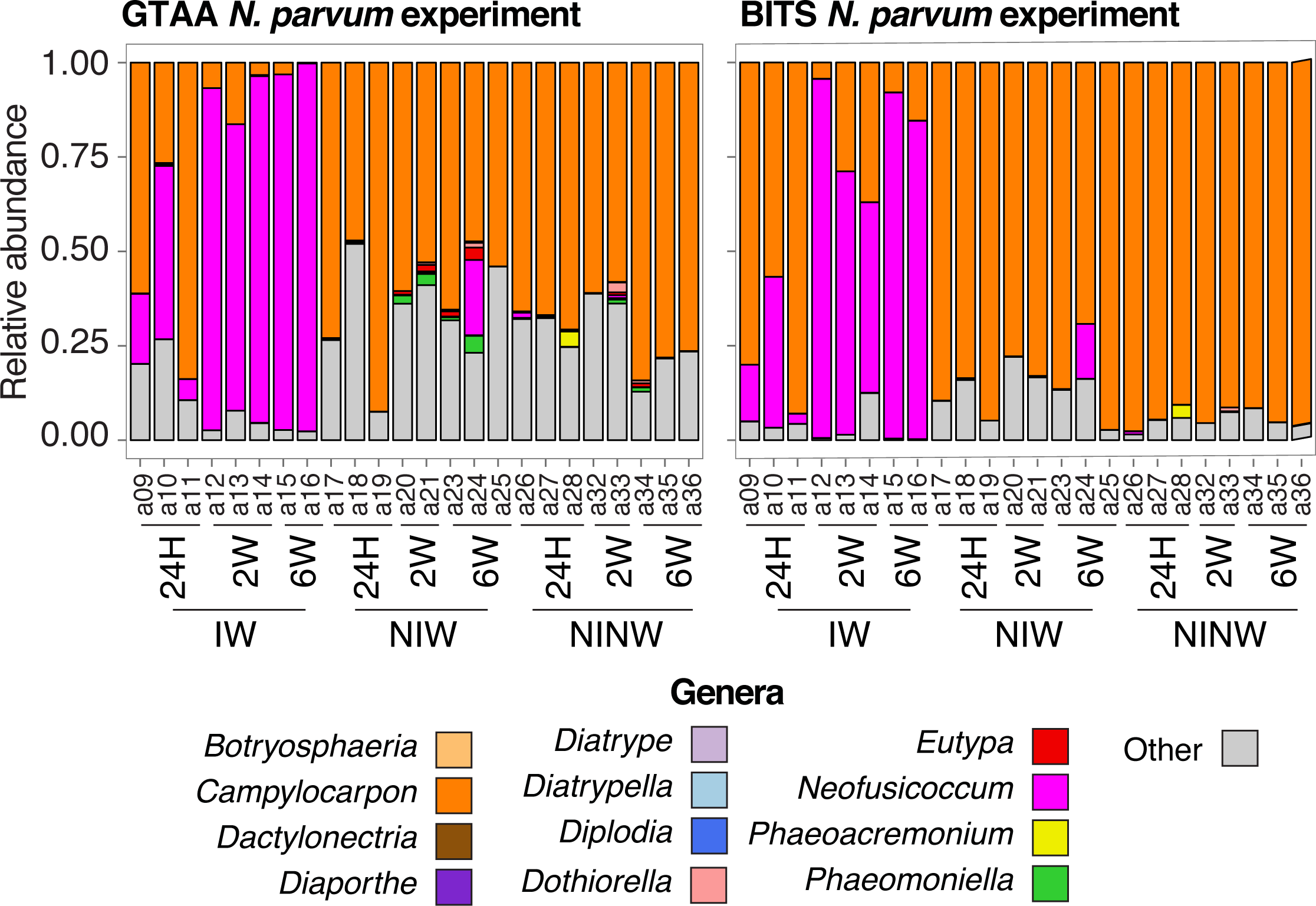
Results of DNA metabarcoding of an infection time-course. Stacked barplots show the relative abundance of the genera detected in a time course experiment after inoculation with *N. parvum.* IW: wound-inoculated with *N. parvum;* NIW: non-inoculated non-wounded; NINW: non-inoculated non-wounded controls.

### Analysis of field samples and comparison with reference-based shotgun metagenome sequencing

We then tested the primers on naturally infected grapevines. We used the same 28 field samples described in [13], which allowed us to compare the metabarcoding approach with the quantitative taxonomic profiles obtained by a reference-based shotgun metagenome sequencing. The samples were grouped according to symptoms into Eutypa dieback (ED), Esca (ES), wood canker without foliar symptoms (WC), and apparently-healthy (AH). All 28 field samples were amplified with both GTAA and BITS primers, with SP primers used for a subset. Taxonomy assignment based on amplicon metabarcoding detected 14 genera, in addition to those with genomes in the multispecies reference, with abundances > 0.05% in one or more samples (**Additional File 3**). Both GTAA and BITS primer sets identified *Alternaria, Cyphellophora*, and *Penicillium*, whereas *Cladosporium, Aureobasidium, Gibberella*, and *Cryptovalsa* were only identified by GTAA primers, and *Angustimassarina, Exophiala, Erysiphe, Meyerozyma, Acremonium*, and *Vishniacozyma* by BITS primers. GTAA primers revealed species abundances at very similar levels to those obtained by metagenomics analysis (**Figure 5A**), with a strong linear correlation between the two approaches (R = 0.95; **Figure 5B**), which was higher than those of both BITS (R = 0.63) and SP primers (R = 0.27). In agreement with the other results described above, BITS primers underestimated *Eutypa* in Eutypa-dieback samples and overrepresented *Phaeomoniella* in Esca samples. The even weaker correlation obtained with SP primers was due to the strong bias against *Eutypa* and *Diaporthe.* GTAA primers showed stronger correlations across all genera of trunk pathogens (0.89 < *R <* 0.99, **Figure 5C**) compared to those of BITS (0.58 < *R* < 0.75). Both primer sets showed a low correlation for *Neofusicoccum*, likely due to the low abundance of this genus in the samples assayed. Overall, our findings confirm the universal primers have a significant bias against important taxa and were outperformed by our GTAA primers for trunk pathogens.

**Figure 5.**
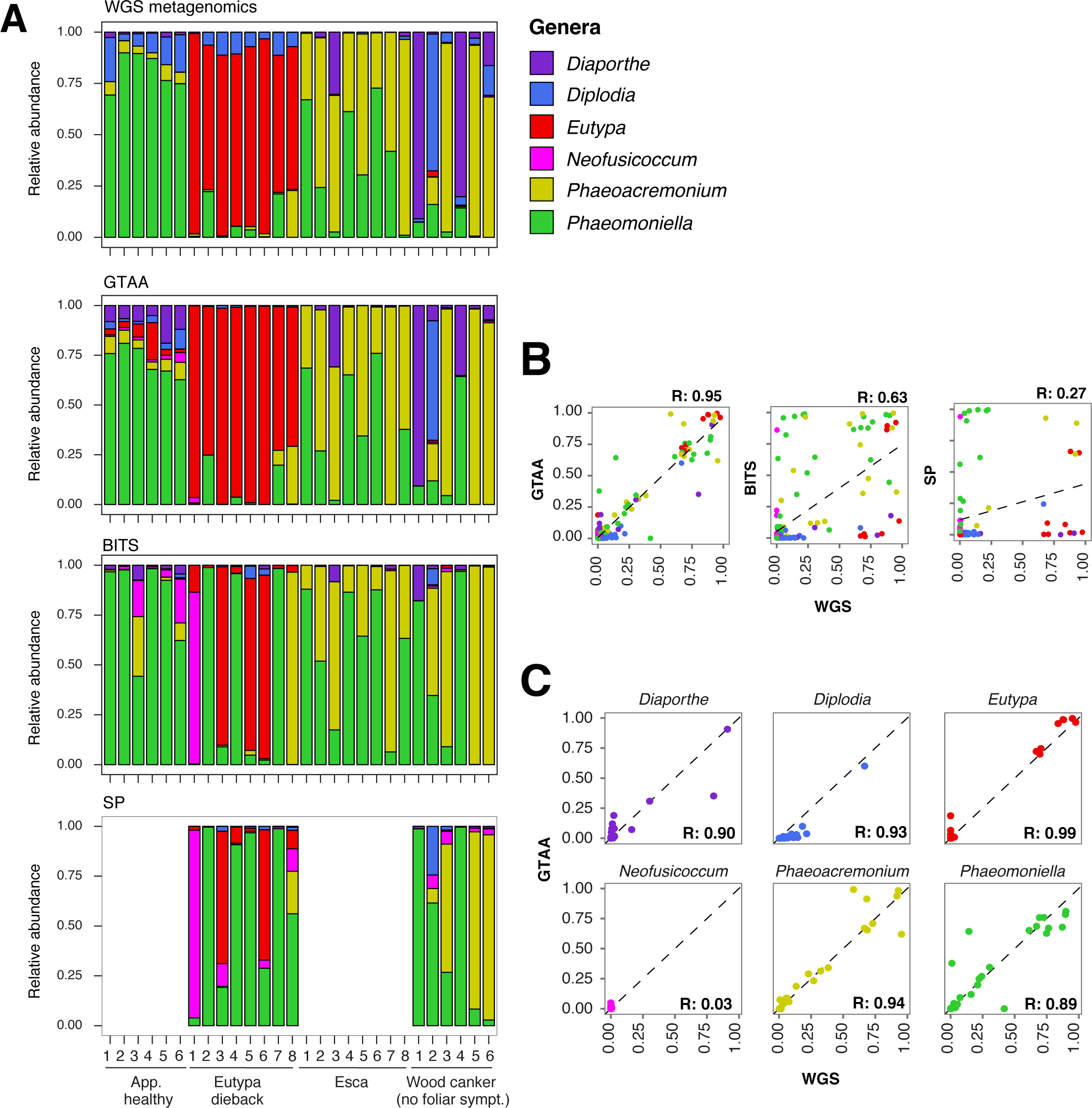
Results of DNA metabarcoding of field samples and comparisons with shotgun whole-genome metagenomics (WGS). (A) Stacked barplots showing the relative abundance of the genera detected by WGS and DNA metabarcoding with GTAA, BITS, and SP primers. (B) Scatter plots showing the correlations of the relative abundance obtained by DNA metabarcoding and WGS. (C) Scatter plots showing the correlations of the relative abundance separately for each genus obtained by DNA metabarcoding using the GTAA primers and WGS.

## DISCUSSION

In this study, we tested the application of DNA metabarcoding to profile the fungal taxa associated with grapevine trunk diseases. We show that DNA metabarcoding of ribosomal ITS amplified with commonly-adopted universal primers consistently misrepresented the abundance of important trunk pathogen species, such as *Eutypa* and *Phaeomoniella*. The customization of primer design using trunk pathogen sequences as template led to improved the results with greater sensitivity. This was likely due to greater homology between the GTAA primers and the ITS of the grapevine trunk pathogens they target. On average the sequence identity of the grapevine trunk pathogen targets was significantly greater with the GTAA primers (97.4 ± 5.5%; *P* < 2e-16) than with the other universal primers (BITS: 90.2 ± 7.1%; SP: 83.3 ± 0.2.2%) used in the study. Amplification bias of universal ITS primers due to higher levels of mismatches for certain taxonomic group were observed previously using *in silico* PCR [54, 55]. Importantly, we also showed that the GTAA primers had higher sensitivity while maintaining a precision threshold for taxonomic assignment of 97%, suggesting that the customization of the target region also played a role in improving the DNA metabarcoding for these organisms. We should stress out that BITS and SP primers are not the only available universal primers and the goal of this study was not to provide a comprehensive survey of all universal ITS metabarcoding primers. BITS and SP were selected, because they are both widely used DNA barcoding primers, including in studies conducted on vineyard and wine must samples <u>[46-51]</u>. We cannot rule out that other universal primers that were not tested in this study may have performed differently. However, the results presented in this study show that universal primers may not be always appropriate to study a fungal community and, when fungal community composition is available, researchers should consider customizing their DNA metabarcoding primers. In addition, we illustrate the value of assessing both the amplification and taxonomy usefulness of the metabarcoding primers *in silico* prior to downstream wet lab evaluations.

In addition to customization of primers, the inclusion of other DNA barcodes should help overcome some of the limitations associated with the ITS region, such as copy number variation between and within species and low resolution in separating some phylogenetically closely related fungal species [42, 56]. For example, the ITS region does not accurately identify species of plant-pathogenic fungi like *Alternaria, Botryosphaeria*, and *Diaporthe* [57]. The genera for which the GTAA primers consistently underestimated abundances like *Lasiodiplodia, Botryosphaeria, Diplodia*, and *Diaporthe* are known to be difficult to be resolved with the ITS region alone [57]. The high correlation between metagenomics and metabarcoding results using the GTAA primers suggest that copy number variation of the ITS region is not an overwhelming issue for the grapevine trunk pathogens present in the field samples. Nonetheless, we expect that the inclusion of additional barcodes, such as β-tubulin and elongation factor 1-α, will help increase accuracy of taxonomic identification at the species level and help measure those genera for which the ITS is known not to be effective [20, 23, 58, 59].

## CONCLUSIONS

As trunk diseases are complex diseases caused by mixed infections, DNA metabarcoding should provide a rapid and effective method for high-throughput multispecies identification overcoming the limitations of currently applied diagnostic methods. Universal primers are advantageous in exploratory analysis where *a priori* knowledge on the taxonomic composition of the samples is limited or not available. However, a more targeted approach should be used when the objective is to study a more defined group of microorganisms, like the grapevine trunk pathogens which symptoms have been consistently associated with certain fungal species [4, 20, 60]. Overall, the results presented here demonstrated that DNA metabarcoding can be applied to grapevine trunk diseases. With further improvement of taxonomic identification by combining multiple barcoding loci and of quantification by measurement of direct correlation between fungal biomass and PCR amplification cycles, we envision DNA metabarcoding to be routinely applied in trunk pathogen research and diagnostics. DNA metabarcoding provides multiple advantages to methods employed in the past. Namely, there is no need of fungal isolation, it allows high number of samples to be analyzed at the same time given the multiplexing potential of the technology, and takes advantage of the constantly improving high-throughput sequencing technologies. Since wood pathogens may remain asymptomatic in young, non-stressed vines, propagation material may contain latent fungal infections and may become symptomatic after planting and serve as a source of inoculum for further infections of potentially clean plants. Methods of virus detection and eradication have been crucial in ensuring that the material in germplasm repositories and clean plant programs is free of known viruses. By allowing the rapid testing of large number of wood samples from mother plants in foundation blocks and propagation material in nurseries, we expect that the applications of metabarcoding to trunk pathogen diagnostics will help reduce the amount of trunk pathogens introduced into vineyards at planting as well as the incidence of young vine decline. Our results also demonstrated that primer customization and testing are crucial to ensure the validity of DNA metabarcoding results.

## METHODS

### Metabarcoding primers targeting grapevine trunk-associated Ascomycetes (GTAA)

Ribosomal Internal transcribed spacer (ITS) sequences of trunk pathogens and other wood-colonizing fungi of grape, specifically in the Division Ascomycota, were retrieved from the TrunkDiseaseID.org database <u>[20]</u>. Sequences were aligned using ClustalW2 (v2.1; <u>[61]</u>) to identify conserved regions. Sequence alignment was used as input for the metabarcoding primer design software Primer Prospector v1.0.1 [62], using a sensitivity threshold of 80% and an initial primer seed size of 5 bp. The ITS sequence of *E. lata* (GeneBank KU721859.1) was used as a ‘reference’. Primers were selected based on median amplicon size, and mismatches, gaps, and numbers of matches to the sequences in the database. The base pairs ‘AG’ were used as a linker between the primer and an eight-nucleotide barcodes on the 5’ region of the forward primer sequence. Barcode sequences were as described in [63]. A list of barcoded forward GTAA primers is listed in **Additional File 4.**

A custom database was compiled with full length ITS sequences of species in the following genera: *Botryosphaeria, Diplodia, Dothiorella, Lasiodiplodia, Neofusicoccum, Phaeomoniella, Diaporthe, Phaeoacremonium, Diatrype, Diatrypella, Eutypa, Xylaria, Cylindrocarpon, Campylocarpon, Dactylonectria, Ilyonectria*, and *Neonectria.* Sequences were retrieved from the NCBI GenBank repository. Completeness of the ITS sequences was validated using the hidden Markov models-based software ITSx <u>[64]</u>. Only sequences spanning the entire ITS region (ITS1, 5.8S, and ITS2) were kept for downstream analysis. Species and GenBank accessions of the complete ITS sequences included in the custom database are listed in **Additional File 2**. To reduce redundancy and identify outliers, the complete ITS sequences were clustered using the UCLUST algorithm <u>[65]</u> integrated in Qiime (v1.9.1; <u>[66]</u>) with 97% identity. The longest representative sequence of each cluster was selected, using the Qiime ‘pick_rep_set.py’ function. All representative sequences were aligned using Mafft v7.271 [67]) with the ‘--auto’ argument and 1,000 iterations. Sequences clustering outside the expected family were removed from the final custom database.

The program Degenerate In-Silico PCR (dispr, https://github.com/douglasgscofield/dispr) was used to predict and evaluate the amplification of sequences of the custom ITS database, using our GTAA primers, and universal BITS [45], and SP <u>[46]</u> primers. Dispr allowed an amplicon size of 100 to 400 bp, all combinations up to five mismatches in the head of the primer (‘H’ or 5’-most region), and all combinations up to two mismatches in the tail of the primer (‘T’ or the remaining 3’-portion of the primer). The resulting amplicons produced *in silico* were then used for taxonomy assignment with 80% confidence, using Mothur (v1.39.5; [53.]), as it is integrated in Qiime (v1.9.1). The UNITE database v7.2 <u>[68]</u> was used as taxonomic reference. True positives were defined as sequences that were assigned to the expected genus, false positives were sequences assigned to a different genus, and false negatives were sequences not assigned to any genus or were not amplified by dispr.

To generate mock communities, we combined (i) DNA from a healthy grapevine with DNA from pure cultures of three trunk pathogens at different concentrations, or (ii) equal concentrations of DNA from pure cultures of six trunk pathogens. For the former, grape and fungal DNA were combined as follows: 90% grape with 10% *E. lata* isolate Napa209 [69], *Phaeoa. minimum* isolate 1119 [70], or *Phaeom. chlamydospora* isolate C42 [71]; 80% grape with 10% of each of two fungal isolates (in all three pair-wise combinations of the three isolates); and 70% grape DNA with 10% of each of the three fungal isolates. For the latter, equal concentrations of DNA were combined from the same three trunk pathogens and three additional species: *N. parvum* isolate UCD646So [11], *Dia. ampelina* isolate Wolf911 [72], and *Diplodia seriata* isolate SBen831 <u>[73]</u>. Grape DNA was extracted from the leaves of a non-inoculated, non-wounded plant; this DNA template came from a previous experiment <u>[74]</u>. The mock communities containing grape DNA were amplified and sequenced independently three times, whereas the mock community of six fungal DNAs was amplified and sequenced independently five times. Prior to DNA extraction on Potato Dextrose Agar (PDA; Difco laboratories, Detroit, MI). DNA was extracted as described in [73] and measured by Qubit (Life technologies).

To test *in planta* detection of a trunk pathogen at variable levels of infection (i.e., from low to high concentrations of fungal biomass over time), DNAs for the infection time course of *N. parvum* were extracted from the same samples described in <u>[74]</u>. Briefly, 1-year-old potted *V. vinifera* ‘Cabernet Sauvignon’ FPS 19 plants were inoculated with isolate UCD646So mycelia. Woody stems were collected at seven time points: 0 hpi, 3 hpi, 24 hpi, 2 wpi, 6 wpi, 8 wpi, and 12 wpi. Wood samples from 1 cm below the inoculation site were collected using flame-sterilized forceps and immediately placed in liquid nitrogen for nucleic acids extraction. Infections were confirmed by positive recovery of the pathogen after 5-day growth on PDA.

To test *in planta* detection of multiple trunk pathogens in mixed infection (i.e., to characterize the species composition of a naturally established trunk-pathogen community), DNA from the same 28 field samples described in [13] was used to make cross-technology comparisons. These field samples were collected from mature vines (> 8 years-old) showing a variety of the most common symptoms associated with trunk diseases. Wood samples were collected from distinct plants with the following combinations of symptoms: Eutypa dieback foliar and wood symptoms, Esca foliar and wood symptoms, wood symptoms and no foliar symptoms, and apparently healthy plants with no foliar or wood symptoms.

### High throughput sequencing libraries

Each sample was amplified using the unique 8-nt barcode forward primer sets for GTAA and BITS, to enable sample multiplexing. The 25-μl PCR reaction mix contained 2 ng of DNA template, 1X Colorless GoTaq flexi buffer (Promega Corporation, Madison, WI), 1.5 mM MgCl_2_, 0. 1 mg/ml BSA, 0.2 mM dNTPs, 0.4 μiM of each primer, and 1.25 units of GoTaq Flexi DNA polymerase (Promega Corporation, Madison, WI). PCR program (Veriti thermal cycler, Applied Biosystems) was as follows: initial denaturation at 94°C for 3 min, followed by 35 cycles at 94°C for 45 seconds, 55°C for 1 min., and 72°C for 1 min., and a final extension at 72°C for 10 min. In the experiments to assess primer affinity, reactions were stopped after 5, 10, 15, 20, 25, 30, and 35 cycles. Following PCR, amplicon size and uniqueness were verified using gel electrophoresis, and bands were cleaned using Ampure XP magnetic beads (Agencourt, Beckman Coulter). DNA concentration was determined for each purified amplicon using Qubit (Life technologies). For the single isolate validation, amplicons were sequenced with Sanger (DNA Sequencing Facility, University of California, Davis).

For high-throughput sequencing, equimolar amounts of all barcoded amplicons were pooled into a single sample, the total concentration of which was determined by Qubit. Five hundred nanograms of pooled DNA were then end-repaired, A-tailed and single-index adapter ligated (Kapa LTP library prep kit, Kapa Biosystems). After adapter ligation, the sample was size-selected with two consecutive 1X bead-based cleanups; concentration and size distribution were determined with Qubit and Bioanalyzer (Agilent Technologies), respectively. DNA libraries were submitted for sequencing in 250-bp paired-end mode on an Illumina MiSeq (UCDavis Genome Center DNA technologies Core). All FASTQ files with the amplicon sequences separated by barcode were deposited in the NCBI Sequence Read Archive (BioProject: PRJNA485180; SRA accession: SRP156804).

### Amplicon sequencing community analysis

Adapter-trimming was carried out using BBDuk (BBMap v.35.82; http://jgi.doe.gov/data-and-tools/bb-tools/) in paired-end mode with sequence “AGATCGGAAG” and the following parameters: ktrim=r, k=10, mink=6, edist=2, ordered=t, qtrim=f and minlen=150. Adapter-trimmed FASTQ files were then quality-filtered using Trimmomatic v0.36 [75] with paired-end mode, phred33, a sliding windows of 4:19, and a minimum length of 150 bp. Sequencing data were then processed in the Qiime environment v1.9.1 <u>[66]</u>. Barcodes were extracted from the FASTQ files using the “extract_barcodes.py” function with the “-a” argument that attempts read orientation and a barcode length of eight base pairs. The resulting sequences and barcodes were used to tag the reads with “split_libraries_fastq.py”, a threshold quality score of 20, and a barcode size of eight basepairs. Operational taxonomic units (OTUs) were identified with a 99% similarity threshold using the UCLUST algorithm <u>[65]</u> with the reverse strand match enabled (“-z”), and the longest sequence of each OUT was chosen as representative sequence. Taxonomy assignment was carried out using Mothur (v1.39.5; <u>[53]</u> with the UNITE database v7.2 <u>[68]</u> as reference and a 80% confidence threshold. For each sample, sequences were randomly sampled with the function “single_rarefaction.py” from the OTU tables to obtained a total number of sequences per sample equal to the lowest number of reads across GTAA, BITS, and SP datasets. Taxonomy tables at the genus level were then created using “summarize_taxa.py”.

## DECLARATIONS

### Ethics approval and consent to participate

Not applicable

### Consent for publication

Not applicable

### Availability of data and material

All FASTQ files with the amplicon sequences separated by barcode were deposited in the NCBI Sequence Read Archive (BioProject: PRJNA485180; SRA accession: SRP156804).

## Authors’ contributions

Conceived and designed the experiments: AMC and DC. Performed the lab experiments: RFB, AMC, ET, and JD. Performed field sampling: PER. Performed bioinformatic analysis: AMC. Wrote the manuscript: AMC, KB, and DC. All authors read and approved the final manuscript.

## ADDITIONAL FILES

### Additional File 1: Text S1 (FASTA)

Assembled amplicon sequences produced by the GTAA primers and sequenced with Sanger.

### Additional File 2: Table S1 (XLSX)

List of filtered sequences retrieved from the NCBI used as a database for primer testing and test results per primer. TP: True Positive, FP: False Positive and FN: False Negative.

### Additional File 3: Figure S1 (PDF)

Venn diagram of genera detected by the GTAA and BITS primers from the field samples detected with more than 0.05% abundance per sample.

### Additional File 4: Table S2 (XLSX)

List of forward GTAA primers with linker and barcodes

## REFERENCES

1. Munkvold G, Duthie J, Marois J: Reductions in yield and vegetative growth of grapevines due to Eutypa dieback. Phytopathology 1994, 84(2):186–192.

2. Siebert J: Economic impact of Eutypa on the California Wine Grape Industry. Report prepared for the American Vineyard Foundation 2000.

3. Kaplan J, Travadon R, Cooper M, Hillis V, Lubell M, Baumgartner K: Identifying economic hurdles to early adoption of preventative practices: the case of trunk diseases in California winegrape vineyards. Wine Economics and Policy 2016, 5(2):127–141.

4. Bertsch C, RaMírez-Suero M, Magnin-Robert M, Larignon P, Chong J, Abou-Mansour E, Spagnolo A, Clément C, Fontaine F: Grapevine trunk diseases: complex and still poorly understood. Plant Pathology 2013, 62(2):243–265.

5. Whitelaw-Weckert M, Rahman L, Appleby L, Hall A, Clark A, Waite H, Hardie W: Co-infection by Botryosphaeriaceae and Ilyonectria spp. fungi during propagation causes decline of young grafted grapevines. Plant Pathology 2013, 62(6):1226–1237.

6. Graniti A, Bruno G, Sparapano L: Three-year observation of grapevines cross-inoculated with esca-associated fungi. Phytopathologia Mediterranea 2001, 40(3):376–386.

7. Pierron RJ, Pouzoulet J, Couderc C, Judic E, Compant S, Jacques A: Variations in early response of grapevine wood depending on wound and inoculation combinations with Phaeoacremonium aleophilum and Phaeomoniella chlamydospora. Frontiers in plant science 2016, 7:268.

8. Cabral A, Rego C, Nascimento T, Oliveira H, Groenewald JZ, Crous PW: Multi-gene analysis and morphology reveal novel Ilyonectria species associated with black foot disease of grapevines. Fungal Biology 2012, 116(1):62–80.

9. Halleen F, Fourie PH, Crous PW: A review of black foot disease of grapevine. Phytopathologia Mediterranea 2006, 45(4):55–67.

10. Moller W, Kasimatis A, Kissler J: A dying arm disease of grape in California [Eutypa armeniacae]. Plant Disease Reporter 1974.

11. Úrbez-Torres J, Leavitt G, Voegel T, Gubler W: Identification and distribution of Botryosphaeria spp. associated with grapevine cankers in California. Plant Disease 2006, 90(12):1490–1503.

12. Lamichhane JR, Venturi V: Synergisms between microbial pathogens in plant disease complexes: a growing trend. Frontiers in Plant Science 2015, 6(385).

13. Morales-Cruz A, Allenbeck G, Figueroa-Balderas R, Ashworth VE, Lawrence DP, Travadon R, Smith RJ, Baumgartner K, Rolshausen PE, Cantu D: Closed-reference metatranscriptomics enables in planta profiling of putative virulence activities in the grapevine trunk disease complex. Molecular plant pathology 2018, 19(2):490–503.

14. Gramaje D, Armengol J: Fungal trunk pathogens in the grapevine propagation process: potential inoculum sources, detection, identification, and management strategies. Plant Disease 2011, 95(9):1040–1055.

15. Fourie P, Halleen F: Occurrence of grapevine trunk disease pathogens in rootstock mother plants in South Africa. Australasian Plant Pathology 2004, 33(2):313–315.

16. Halleen F, Crous R, Petrin O: Fungi associated with healthy grapevine cuttings in nurseries, with special reference to pathogens involved in the decline of young vines. Australasian Plant Pathology 2003, 32(1):47–52.

17. Sosnowski MR, Wicks TJ, Scott ES: Control of Eutypa dieback in grapevines using remedial surgery. Phytopathologia Mediterranea 2011, 50(4):277–284.

18. Rolshausen PE, Úrbez-Torres JR, Rooney-Latham S, Eskalen A, Smith RJ, Gubler WD: Evaluation of pruning wound susceptibility and protection against fungi associated with grapevine trunk diseases. American Journal of Enology and Viticulture 2010, 61(1):113–119.

19. Gramaje D, Úrbez-Torres JR, Sosnowski MR: Managing Grapevine Trunk Diseases With Respect to Etiology and Epidemiology: Current Strategies and Future Prospects. Plant Disease 2018, 102(1):12–39.

20. Lawrence DP, Travadon R, Nita M, Baumgartner K: TrunkDiseaseID. org: A molecular database for fast and accurate identification of fungi commonly isolated from grapevine wood. Crop Protection 2017, 102:110–117.

21. Pouzoulet J, Mailhac N, Couderc C, Besson X, Daydé J, Lummerzheim M, Jacques A: A method to detect and quantify Phaeomoniella chlamydospora and Phaeoacremonium aleophilum DNA in grapevine-wood samples. Applied microbiology and biotechnology 2013, 97(23):10163–10175.

22. Pouzoulet J, Rolshausen PE, Schiavon M, Bol S, Travadon R, Lawrence DP, Baumgartner K, Ashworth VE, Comont G, Corio-Costet M-F: A method to detect and quantify Eutypa lata and Diplodia seriata-complex DNA in grapevine pruning wounds. Plant disease 2017, 101(8):1470–1480.

23. Úrbez-Torres J, Haag P, Bowen P, Lowery T, O’Gorman D: Development of a DNA macroarray for the detection and identification of fungal pathogens causing decline of young grapevines. Phytopathology 2015, 105(10):1373–1388.

24. Martín MT, Cobos R, Martín L, López-Enríquez L: Real-time PCR detection of Phaeomoniella chlamydospora and Phaeoacremonium aleophilum. Applied and environmental microbiology 2012, 78(11):3985–3991.

25. Lawrence DP, Travadon R, Baumgartner K: Novel Seimatosporium Species from Grapevine in Northern California and Their Interactions with Fungal Pathogens Involved in the Trunk-Disease Complex. Plant Disease 2018, 102(6):1081–1092.

26. Pancher M, Ceol M, Corneo PE, Longa CMO, Yousaf S, Pertot I, Campisano A: Fungal endophytic communities in grapevines (Vitis vinifera L.) respond to crop management. Applied and environmental microbiology 2012:AEM. 07655-07611.

27. Bruez E, Vallance J, Gerbore J, Lecomte P, Da Costa J-P, Guerin-Dubrana L, Rey P: Analyses of the temporal dynamics of fungal communities colonizing the healthy wood tissues of esca leaf-symptomatic and asymptomatic vines. PLOS One 2014, 9(5):e95928.

28. Bruez E, Baumgartner K, Bastien S, Travadon R, Guérin-Dubrana L, Rey P: Various fungal communities colonise the functional wood tissues of old grapevines externally free from grapevine trunk disease symptoms. Australian journal of grape and wine research 2016, 22(2):288–295.

29. Schmidt P-A, Bálint M, Greshake B, Bandow C, Römbke J, Schmitt I: Illumina metabarcoding of a soil fungal community. Soil Biology and Biochemistry 2013, 65:128–132.

30. Whipps J, Hand P, Pink D, Bending GD: Phyllosphere microbiology with special reference to diversity and plant genotype. Journal of applied microbiology 2008, 105(6):1744–1755.

31. Fierer N, Leff JW, Adams BJ, Nielsen UN, Bates ST, Lauber CL, Owens S, Gilbert JA, Wall DH, Caporaso JG: Cross-biome metagenomic analyses of soil microbial communities and their functional attributes. Proceedings of the National Academy of Sciences 2012, 109(52):21390–21395.

32. Prober SM, Leff JW, Bates ST, Borer ET, Firn J, Harpole WS, Lind EM, Seabloom EW, Adler PB, Bakker JD: Plant diversity predicts beta but not alpha diversity of soil microbes across grasslands worldwide. Ecology letters 2015, 18(1):85–95.

33. Xu X, Passey T, Wei F, Saville R, Harrison RJ: Amplicon-based metagenomics identified candidate organisms in soils that caused yield decline in strawberry. Horticulture research 2015, 2:15022.

34. Abdelfattah A, Cacciola SO, Mosca S, Zappia R, Schena L: Analysis of the fungal diversity in citrus leaves with greasy spot disease symptoms. Microbial ecology 2017, 73(3):739–749.

35. Deyett E, Roper MC, Ruegger P, Yang J-I, Borneman J, Rolshausen PE: Microbial Landscape of the Grapevine Endosphere in the Context of Pierce’s Disease. Phytobiomes 2017, 1(3):138–149.

36. Malacrinò A, Schena L, Campolo O, Laudani F, Mosca S, Giunti G, Strano CP, Palmeri V: A metabarcoding survey on the fungal microbiota associated to the olive fruit fly. Microbial ecology 2017, 73(3):677–684.

37. Quaedvlieg W, Groenewald J, de Jesús Yáñez-Morales M, Crous P: DNA barcoding of Mycosphaerella species of quarantine importance to Europe. Persoonia: Molecular Phylogeny and Evolution of Fungi 2012, 29:101.

38. Bulman S, McDougal R, Hill K, Lear G: Opportunities and limitations for DNA metabarcoding in Australasian plant-pathogen biosecurity. Australasian Plant Pathology 2018:1–8.

39. Hodgetts J, Ostojá-Starzewski JC, Prior T, Lawson R, Hall J, Boonham N: DNA barcoding for biosecurity: case studies from the UK plant protection program. Genome 2016, 59(11):1033–1048.

40. Taberlet P, Coissac E, Pompanon F, Brochmann C, Willerslev E: Towards next-generation biodiversity assessment using DNA metabarcoding. Molecular ecology 2012, 21(8):2045–2050.

41. Taberlet P, PruD’homme SM, Campione E, Roy J, Miquel C, Shehzad W, Gielly L, Rioux D, Choler P, Clément J-C: Soil sampling and isolation of extracellular DNA from large amount of starting material suitable for metabarcoding studies. Molecular Ecology 2012, 21(8):1816–1820.

42. Schoch CL, Seifert KA, Huhndorf S, Robert V, Spouge JL, Levesque CA, Chen W, Bolchacova E, Voigt K, Crous PW: Nuclear ribosomal internal transcribed spacer (ITS) region as a universal DNA barcode marker for Fungi. Proceedings of the National Academy of Sciences 2012, 109(16):6241–6246.

43. Bruns TD, White TJ, Taylor JW: Fungal molecular systematics. Annual Review of Ecology and systematics 1991, 22(1):525–564.

44. Coissac E, Riaz T, Puillandre N: Bioinformatic challenges for DNA metabarcoding of plants and animals. Molecular ecology 2012, 21(8):1834–1847.

45. Bokulich NA, Mills DA: Improved selection of internal transcribed spacer-specific primers enables quantitative, ultra-high-throughput profiling of fungal communities. Applied and environmental microbiology 2013, 79(8):2519–2526.

46. Smith DP, Peay KG: Sequence depth, not PCR replication, improves ecological inference from next generation DNA sequencing. PLOS one 2014, 9(2):e90234.

47. Bokulich NA, Thorngate JH, Richardson PM, Mills DA: Microbial biogeography of wine grapes is conditioned by cultivar, vintage, and climate. Proceedings of the National Academy of Sciences 2014, 111(1):E139–E148.

48. Setati ME, Jacobson D, Bauer FF: Sequence-based analysis of the Vitis vinifera L. cv Cabernet Sauvignon grape must mycobiome in three South African vineyards employing distinct agronomic systems. Frontiers in microbiology 2015, 6:1358.

49. Bokulich NA, Ohta M, Richardson PM, Mills DA: Monitoring seasonal changes in winery-resident microbiota. PLOS One 2013, 8(6):e66437.

50. Wang C, García-Fernández D, Mas A, Esteve-Zarzoso B: Fungal diversity in grape must and wine fermentation assessed by massive sequencing, quantitative PCR and DGGE. Frontiers in microbiology 2015, 6:1156.

51. Bokulich NA, Collins TS, Masarweh C, Allen G, Heymann H, Ebeler SE, Mills DA: Associations among wine grape microbiome, metabolome, and fermentation behavior suggest microbial contribution to regional wine characteristics. MBio 2016, 7(3):e00631–00616.

52. Walters W, Hyde ER, Berg-Lyons D, Ackermann G, Humphrey G, Parada A, Gilbert JA, Jansson JK, Caporaso JG, Fuhrman JA: Improved bacterial 16S rRNA gene (V4 and V4-5) and fungal internal transcribed spacer marker gene primers for microbial community surveys. Msystems 2016, 1(1):e00009–00015.

53. Schloss PD, Westcott SL, Ryabin T, Hall JR, Hartmann M, Hollister EB, Lesniewski RA, Oakley BB, Parks DH, Robinson CJ: Introducing mothur: open-source, platform-independent, community-supported software for describing and comparing microbial communities. Applied and environmental microbiology 2009, 75(23):7537– 7541.

54. Bellemain E, Carlsen T, Brochmann C, Coissac E, Taberlet P, Kauserud H: ITS as an environmental DNA barcode for fungi: an in silico approach reveals potential PCR biases. BMC microbiology 2010, 10(1):189.

55. Clarke LJ, Soubrier J, Weyrich LS, Cooper A: Environmental metabarcodes for insects: in silico PCR reveals potential for taxonomic bias. Molecular ecology resources 2014, 14(6):1160–1170.

56. Kiss L: Limits of nuclear ribosomal DNA internal transcribed spacer (ITS) sequences as species barcodes for Fungi. Proceedings of the National Academy of Sciences 2012, 109(27):E1811–E1811.

57. Kashyap PL, Rai P, Kumar S, Chakdar H, Srivastava AK: DNA Barcoding for Diagnosis and Monitoring of Fungal Plant Pathogens. In: Molecular Markers in Mycology. Springer; 2017: 87–122.

58. Travadon R, Lawrence DP, Rooney-Latham S, Gubler WD, Wilcox WF, Rolshausen PE, Baumgartner K: Cadophora species associated with wood-decay of grapevine in North America. Fungal biology 2015, 119(1):53–66.

59. Úrbez-Torres J, Haag P, Bowen P, O’Gorman D: Grapevine trunk diseases in British Columbia: incidence and characterization of the fungal pathogens associated with esca and Petri diseases of grapevine. Plant Disease 2014, 98(4):469–482.

60. Mugnai L, Graniti A, Surico G: Esca (black measles) and brown wood-streaking: two old and elusive diseases of grapevines. Plant disease 1999, 83(5):404–418.

61. Larkin MA, Blackshields G, Brown N, Chenna R, McGettigan PA, McWilliam H, Valentin F, Wallace IM, Wilm A, Lopez R: Clustal W and Clustal X version 2.0. Bioinformatics 2007, 23(21):2947–2948.

62. Walters WA, Caporaso JG, Lauber CL, Berg-Lyons D, Fierer N, Knight R: PrimerProspector: de novo design and taxonomic analysis of barcoded polymerase chain reaction primers. Bioinformatics 2011, 27(8):1159–1161.

63. Harris JK, Sahl JW, Castoe TA, Wagner BD, Pollock DD, Spear JR: Comparison of normalization methods for construction of large, multiplex amplicon pools for next-generation sequencing. Applied and environmental microbiology 2010, 76(12):3863– 3868.

64. Bengtsson-Palme J, Ryberg M, Hartmann M, Branco S, Wang Z, Godhe A, Wit P, Sánchez-García M, Ebersberger I, Sousa F: Improved software detection and extraction of ITS1 and ITS2 from ribosomal ITS sequences of fungi and other eukaryotes for analysis of environmental sequencing data. Methods in ecology and evolution 2013, 4(10):914–919.

65. Edgar RC: Search and clustering orders of magnitude faster than BLAST. Bioinformatics 2010, 26(19):2460–2461.

66. Caporaso JG, Kuczynski J, Stombaugh J, Bittinger K, Bushman FD, Costello EK, Fierer N, Pena AG, Goodrich JK, Gordon JI: QIIME allows analysis of highthroughput community sequencing data. Nature methods 2010, 7(5):335.

67. Katoh K, Standley DM: MAFFT multiple sequence alignment software version 7: improvements in performance and usability. Molecular biology and evolution 2013, 30(4):772–780.

68. UNITE_Community: UNITE QIIME release. In. Edited by Community U, Version 01.12.2017 edn; 2017.

69. Travadon R, Baumgartner K: Molecular polymorphism and phenotypic diversity in the eutypa dieback pathogen eutypa lata. Phytopathology 2015, 105(2):255–264.

70. Massonnet M, Morales-Cruz A, Minio A, Figueroa-Balderas R, Lawrence DP, Travadon R, Rolshausen PE, Baumgartner K, Cantu D: Whole-Genome Resequencing and Pan-Transcriptome Reconstruction Highlight the Impact of Genomic Structural Variation on Secondary Metabolite Gene Clusters in the Grapevine Esca Pathogen Phaeoacremonium minimum. Frontiers in Microbiology 2018, 9(1784).

71. Latham SR: Etiology, epidemiology and pathogen biology of Esca disease of grapevines in California. 2005.

72. Lawrence DP, Travadon R, Baumgartner K: Diversity of Diaporthe species associated with wood cankers of fruit and nut crops in northern California. Mycologia 2015, 107(5):926–940.

73. Morales-Cruz A, Amrine KC, Blanco-Ulate B, Lawrence DP, Travadon R, Rolshausen PE, Baumgartner K, Cantu D: Distinctive expansion of gene families associated with plant cell wall degradation, secondary metabolism, and nutrient uptake in the genomes of grapevine trunk pathogens. BMC genomics 2015, 16(1):469.

74. Massonnet M, Morales-Cruz A, Figueroa-Balderas R, Lawrence DP, Baumgartner K, Cantu D: Condition-dependent co-regulation of genomic clusters of virulence factors in the grapevine trunk pathogen Neofusicoccum parvum. Molecular plant pathology 2018, 19(1):21–34.

75. Bolger AM, Lohse M, Usadel B: Trimmomatic: a flexible trimmer for Illumina sequence data. Bioinformatics 2014, 30(15):2114–2120.

